# Locomotor strategies inside the blind walks: freely moving previsual rat pups in the open field test

**DOI:** 10.1101/2022.02.07.479442

**Authors:** I.S. Midzyanovskaya, V. V. Strelkov

## Abstract

Blind walks of previsual rat pups in the open field test were considered as a supposedly random process. At this age, immature rats should rely only on non-visual modalities for sampling a novel space and constructing putative cognitive maps. Spontaneous locomotion of outbred Wistar rat infants (n=51) on their 13th postnatal day was tracked for 2 minutes and analyzed offline for the presence of random components and/or strategies of locomotion. Three distinct patterns were observed. A large portion of the pups (n=22) performed sub-diffusive localized walks in the center. A smaller cohort (n=9) undertook almost immediate quasi-linear raids in a random direction, thus succeeded in reaching a shelter (i.e., walls’ vicinity). The rest (n=20) demonstrated a mixed strategy: localized walks interspersed with quasi-linear raids. An algorithm for automated segmentation of the open field trajectories into the localized walks and quasi-linear fragments was developed; the fragments were analyzed separately.

Statistical analysis shows the localized walks to be diffusive (i.e., Brownian) only within 2-3 seconds, but essentially sub-diffusive for longer time scales. The autocorrelation function shows that sub-diffusion was caused by the trajectories’ re-attraction. Self-odor traces can be a physical cue ensuring the effect.

Gender or body weights were not significant predictors for any locomotor parameters. The localized walks can be considered as a primary level of idiothetically-cued path integration in previsual pups, whereas the run trials can be a form of an active escape response. The applied method of space potentials avails the automated functional segmentation of locomotor tracks and thus brings advantages of big data analysis for studying non-visual navigation in animals.

## INTRODUCTION

“Blind pup poking” is a literal symbol of random processes. However, nocturnal rodents are often able to navigate through available space without significant visual inputs [Jacobs, 2012]. In immature rat pups active locomotion precedes the ability to use visual landmarks. Olfactory and haptic sensing precedes the first locomotor abilities, which are forelimb-driven propulsion and pivoting. Active whisking starts from PND 8-9, quadrupedal locomotion substitutes forelimb-driven crawling at PND 10-11, digitigrade gait prevail on the planitigrade one since PND 14-15 [Jamon and Clarac,1998; Clarac et al, 2004]. The latter transition happens on the border of eye opening, which is usually observed at PND 14-15. Functionally blind pups are able to return to their nest if drawn away nipples-attached [Renner and Pierre, 1998]. Spatial exploration begins with the onset of spontaneous locomotion from the nest at 10-12 PND [Renner and Pierre, 1998]. Presumably their navigation is guided by the siblings’ odor and sounds. Since the developed locomotor abilities imply a subsequent development of locomotor strategies, the previsual ontogenetic stage might provide insights into nonvisual basis of navigation.

Here we examined locomotor tracks of previsual rat pups in the open field test, in the absence (or insufficiency) of external cues. We selected the short time window (namely PND 13), when rat infants move actively but cannot rely on the visual inputs. At this age, the egocentric navigational system should develop to allocentric one, by including environmental cues perceived by whisking [Wills et al, 2014; Peyrache et al, 2017] and sniffing [Jacobs et al, 2003; Jacobs et al, 2012]. In the absence or insufficiency of tactile and olfactory stimuli, we hope to observe a locomotion with random (no strategy to move) and non-random (one or more strategy to move) components.

Random walk is one of the basic biological models since the discovery of Brownian motion in pollen particles. The Brownian motion implies that each step is made in a completely random direction. It leads to the standard diffusion behaviour, e.g. seen in ink drops diffusing on a water surface. Animals, unlike the idealized physical bodies, are usually incapable of very sharp changes of direction and tend to keep their locomotion “ahead”, i.e. in the caudorostral direction. Thus, a natural correlation between the successive step orientations exists, which is termed ‘persistence’ [Patlak 1953, Codling et al, 2008]. The highest persistence should lead to straightforward locomotion. There is a mechanism to support the locomotor persistency. Namely, a widespread network of the so called head direction neurons starts its maturation at the age of PND 13 [Dudchenko et al, 2019] and is further refined by eye opening. Therefore, straightforward runs and classical random walks are expectable at this age [Tocker et al, 2018]. Several types of tracks’ patterns were indeed seen in experimental data. To differentiate between the smooth (quasi-linear) and entangled (localized walks) patterns, we applied the method of space potentials. The method is widely used for data clusterization in physics but (to our knowledge) has not been applied to behavioural sciences so far.

## METHODS

### Animals

7 litters of normally developing outbred Wistar rats were enrolled in this study. Timely pregnant Wistar dams were kept individually in standard plastic cages, with food and water ad lib, under an artificial 12h light/dark regime (light on at 08/00 a.m). The offspring appearance was monitored daily, the day of birth was assigned as “postnatal day 0” (PND 0). Litters were intact until the day of experiment. On the test day (PND 13), the experimenters watched for the moments when the dams naturally left their nests. At this time point the dams were separated to a transportation cage and the pups were allowed to calm in their huddle for about 30 minutes. Then, the pups were gently grasped individually from the huddle’s periphery, and transported to the experimental chamber next door. To reduce a putative olfactory shock, the nytril gloves used by experimenters were let to adjust to a familiar odour. For this purpose, the gloves had been kept in a closed container with standard pellet foods used for the rats, for 2-3 days prior to the testing. Being grasped, the pup was individually released in the center of an open field arena (black chamber 59cm*59 cm, see Fig. 1), with a camera and a microphone placed 1m above. A standard spatial orientation of the landed pup was the head away from the experimenter, directed to the middle of the opposite wall, see Fig.2 a,b,c). The pup’s placement was a start for video tracking which lasted for 120 seconds. After the test, the pups were sexed, weighed, and numbered (the information is summarized in Table 1), and then returned to their mothers. The arena was cleaned with 50% ethanol solutions and wiped with a clean paper towel after each session.

**Fig.1.**
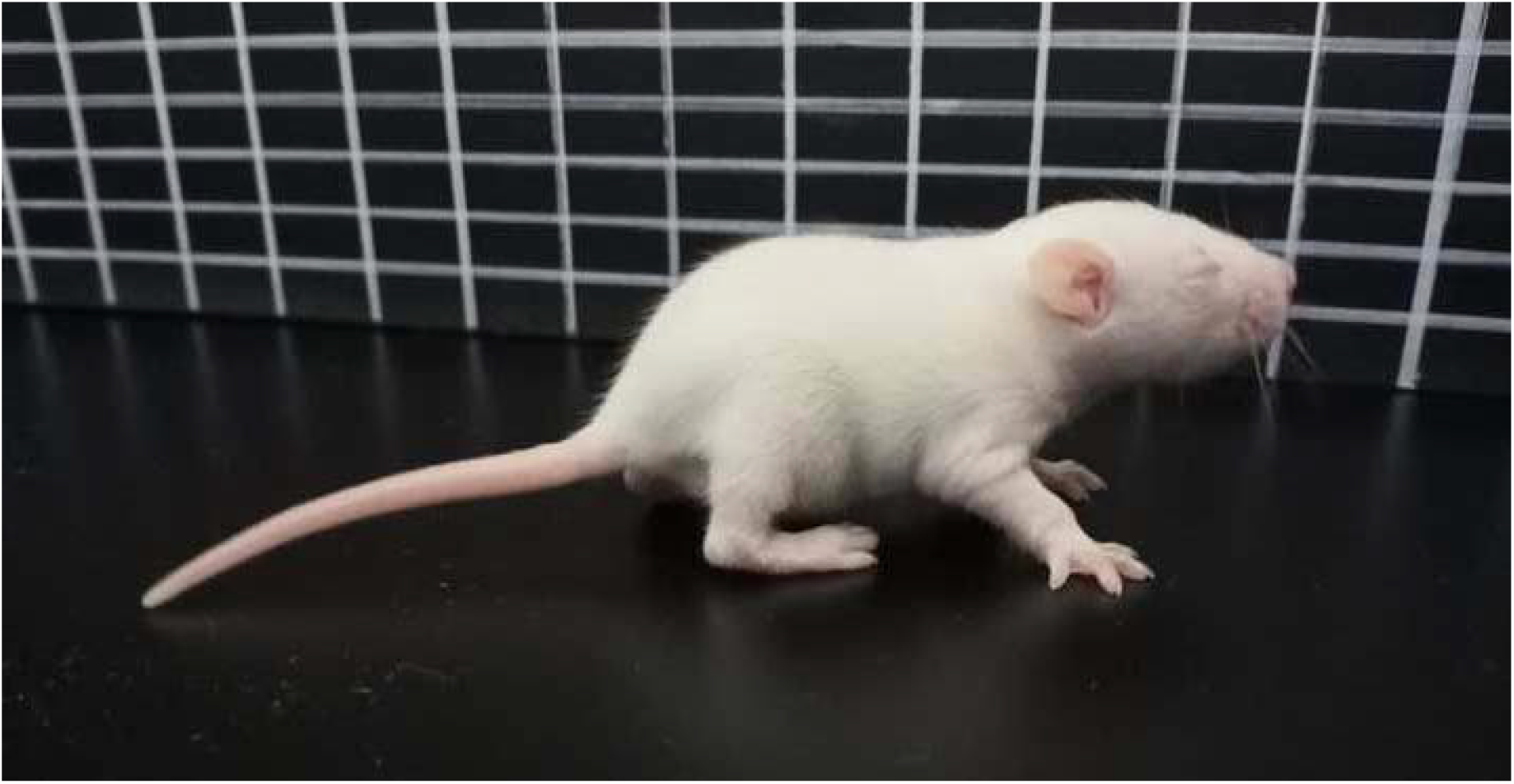
A previsual Wistar rat pup (PND 13) in an open field arena.

**Fig.2.**
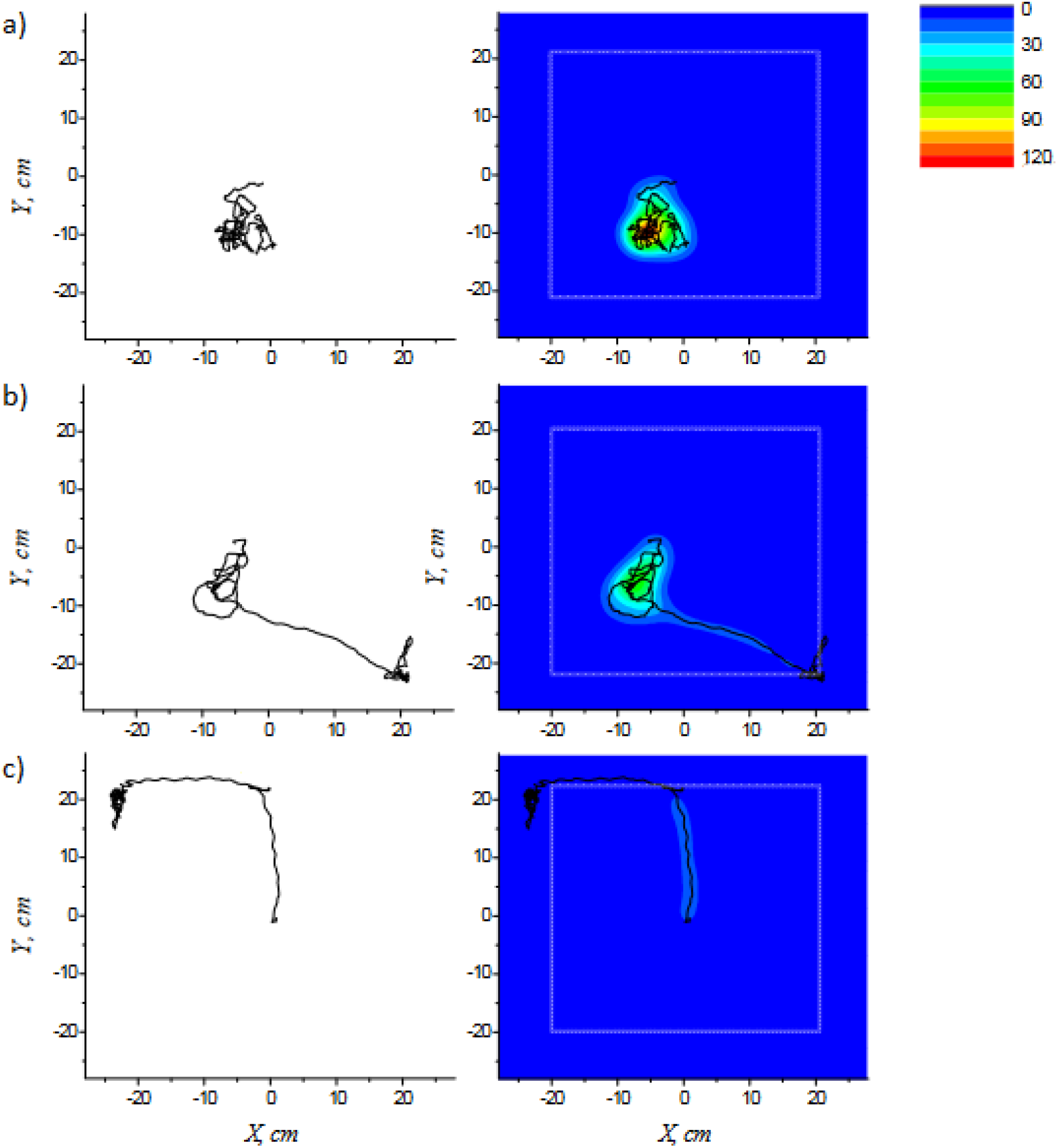
Three examples of trajectories (left panels) segmented into quasi-linear runs and localized walks (right panels) according to the space potential method (see Appendix). Grey hatched lines demark the zone of thigmotactic locomotion. **a.)** An example of the “drift” strategy (group A)’, i.e. localized walks with minimal displacement. **b.)** An example of the “drift and run” strategy (group B), with localized walks interspersed with quasi-linear raids. **c.)** An example of the “run” strategy (group C), a quasi-linear raid ended with thigmotaxis. Light-blue shadings envelope the trajectories’ fragments corresponding to quasi-linear runs, green to red shadings mark those corresponding to localized walks. Thigmotaxis was considered as a special type of locomotion and not analyzed here

**Table 1.**
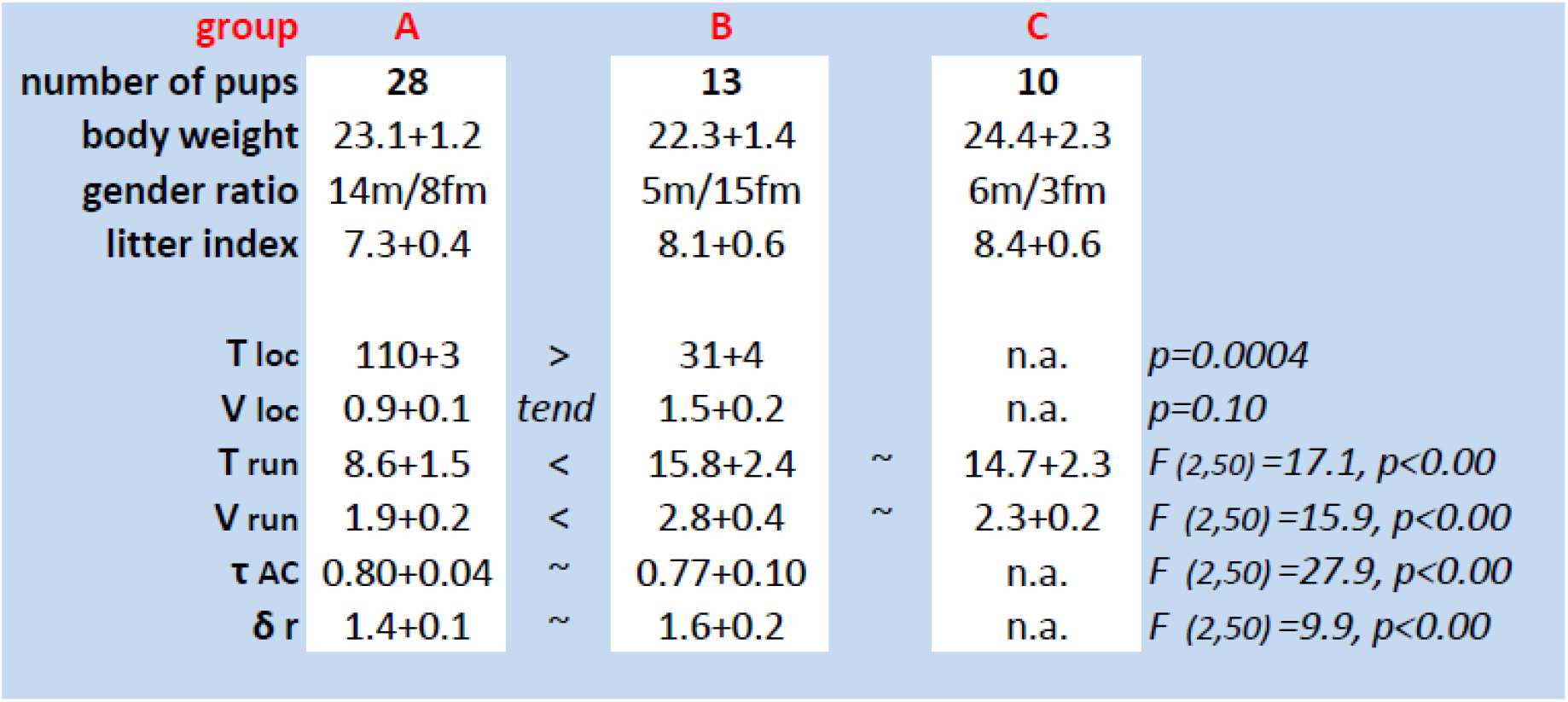
Physiological and locomotor characteristics of the behavioural groups. The first line (n) is for the number of pups in the corresponded group. Mean values and SEM are given for the body weight, gender ratio, litter index, and T_loc_ (duration of localized walks), V_loc_ (velocity of localized walks), T_run_ (duration of quasi-linear runs), V_run_ (velocity of the quasi-linear runs), τ_ac_ (autocorrelation time), and δr (free run path). The statistical significance of the group differences, if any, is given at the right end of each row. The “tend” sign stands for tendency, “n.a” stands for “not applicable”.

| group | A |  | B |  | C |  |
| --- | --- | --- | --- | --- | --- | --- |
| number of pups | 28 |  | 13 |  | 10 |  |
| body weight | 23.1+1.2 |  | 22.3+1.4 |  | 24.4+2.3 |  |
| gender ratio | 14m/8fm |  | 5m/15fm |  | 6m/3fm |  |
| litter index | 7.3+0.4 |  | 8.1+0.6 |  | 8.4+0.6 |  |
| $T_{loc}$ | 110+3 | > | 31+4 | | n.a. | $p=0.0004$ |
| $V_{loc}$ | 0.9+0.1 | tend | 1.5+0.2 | | n.a. | $p=0.10$ |
| $T_{run}$ | 8.6+1.5 | < | 15.8+2.4 | ~ | 14.7+2.3 | $F(2,50)=17.1, p<0.00$ |
| $V_{run}$ | 1.9+0.2 | < | 2.8+0.4 | ~ | 2.3+0.2 | $F(2,50)=15.9, p<0.00$ |
| $\tau_{AC}$ | 0.80+0.04 | ~ | 0.77+0.10 | | n.a. | $F(2,50)=27.9, p<0.00$ |
| $\delta r$ | 1.4+0.1 | ~ | 1.6+0.2 | | n.a. | $F(2,50)=9.9, p<0.00$ |

The experiments were carried out in accordance with the National Institutes of Health Guide for the Care and Use of Laboratory Animals. The experimental protocol was approved by the Institutional Animal Care Committee. All efforts were made to minimize animal discomfort.

### Trajectories analysis

The arena was virtually divided into the “open space” and the “area of thigmotaxis”. The thigmotactic zone covered the area of the walls’ proximity, i.e. of the possible haptic detection of the wall by vibrissae (3cm from the arena bottom’s borders).

### Thigmotactic motion was not considered

The random walks of animals can be affected by geometry of the space and its boundedness [Christensen et al, 2021]. In rodents, the boundary effects are the most prominent, since they prefer to move thigmotactically minimizing the risk of predation and maximizing security [Nemati et al, 2013;Whishaw et al, 2006]. In our experiments can we see it as a phenomenon of obligatory thigmotaxis: a pup keeps moving or sitting in the walls’ vicinity after the first contact (Fig.2B,C). In previsual pups tactile contacts with vertical boundaries additionally allow the brains to correct internal errors of the developing navigation system [Bjerknes et al, 2015]. Therefore, thigmotactic motion was considered as a special type of 1D-locomotion and excluded from our analysis. We extracted only open space fragments of the blind tracks.

### Taxis in the central zone

The tracks were analyzed offline, as described in Appendix. Briefly, the tracks were filtered and smoothed to obviate the possible tracking artifacts (see Appendix sections A I-III for the details). Each point of the track was assigned with a field potential, distributed normally with a width comparable to pups’ body size (see section A IV). The field potentials were summarized for each point and transposed to colour codes for visualization (Fig.2 a,b,c) and trajectories’ segmentation. The points with potential exceeding a certain threshold (green colour and above) were assigned to the localized walk episodes, others (light-blue colour) - to quasi-linear run episodes.

Clearly, the ‘quasi-linear’ trajectories (shown in Fig.2c) contain almost no localized walks. Other tracks (for instance, that shown in (Fig.2 a,b) contain such walks, and below we will study the extent of their randomness.

If the motion is completely random the squared displacement (see the accurate definition in section A VI) during time δ_t_ grows linearly with time. This model is denoted sometimes as a ‘walk of a drunk man’; it describes Brownian motion and many other processes in physics, biology, etc [see Codling et al, 2008 and references therein]. Analyzing the pups’ trajectories we find that the mean squared displacement does not grow linearly with time for the localized walk segments (see Fig. A5, in the Appendix). To study this behaviour in more detail we need to:

- select the localized walk fragments of the tracks and process these fragments
- normalize the results so that averaging over different animals can be valid.

After the trajectories’ segmentation, the longest episodes of localized walks (if any) and quasi-linear runs (if any) for each pup were extracted. We analyzed the time spent in the particular locomotor mode and the velocities of moving along the corresponding segment T_loc_, V_loc_, T_run_, V_run_, respectively. The velocity autocorrelation function was calculated for the localized walks only, since quasi-linear locomotion produces autocorrelation function undistinguishable from unity.

### Clustering the data

The pups were assigned to a group according to their individual values of T_loc_, falling into one of the three chosen ranges (0-10s, 10-50s, 50-120s) in the corresponding histogram (Fig.3 a).

**Fig3.**
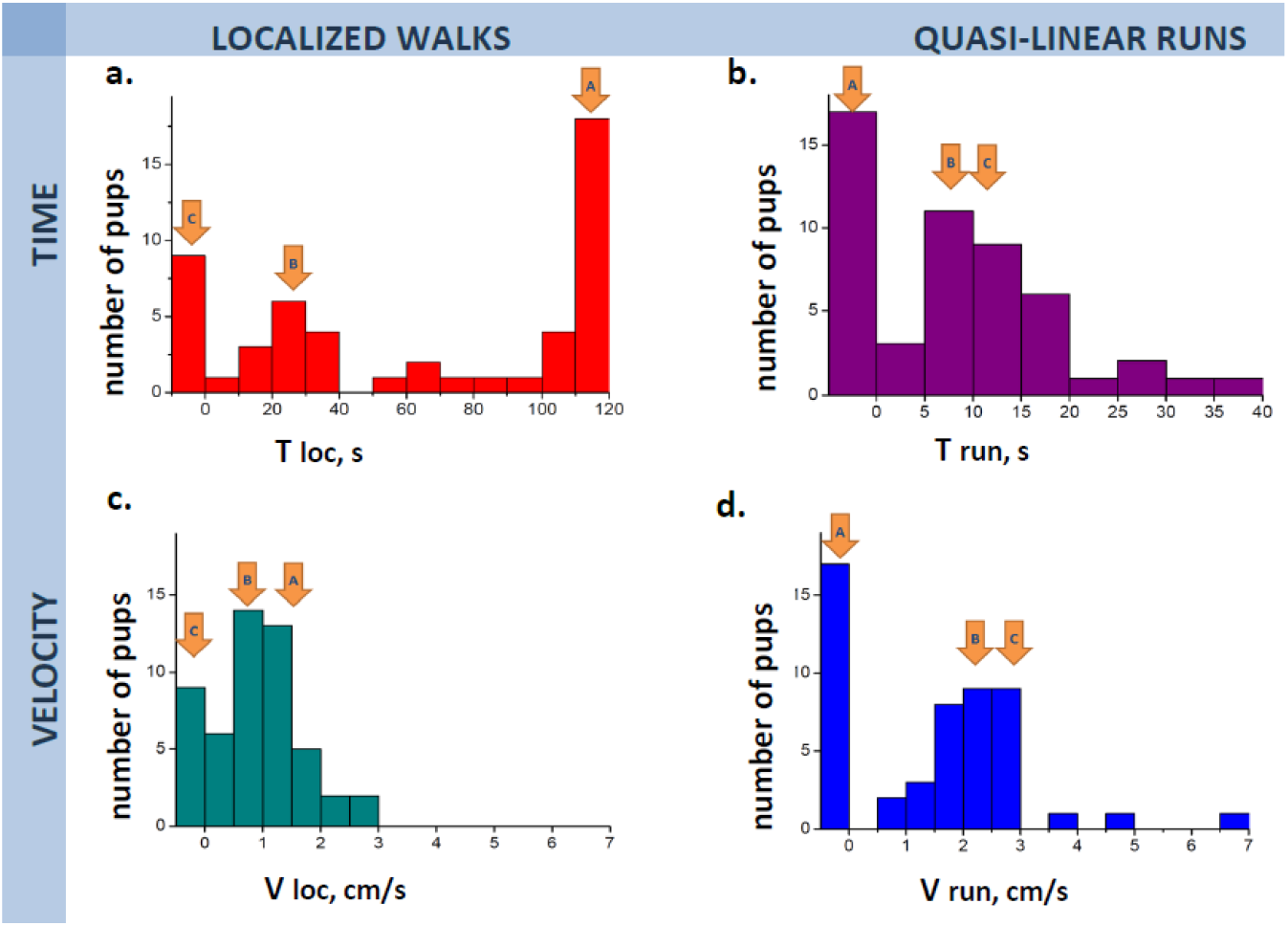
The histograms built for T_loc_ (duration of localized walks), V_loc_(velocity of localized walks), T_run_ (duration of quasi-linear runs), V_run_ (velocity of the quasi-linear runs). The corresponding behavioural groups are marked by orange arrows above the histogram. **a)** The distribution of T_loc_ durations. Note that there are 3 distinct entities: zero-group (no localized walks, group C), moderate T_loc_ (group B) and prolonged T_loc_ (group A). The data are clustered according to the following T_loc_ intervals: 0-10s, 10-50s, and 50-120s. **b)** The distribution of T_run_ durations. Zero peak here corresponds to the behavioural group A (no run trials), group B and C form a single entity. **c)** The distribution of V_loc_ values. The peak at zero V_loc_ here corresponds to the behavioural group C, whereas groups A and B form a single entity. **d)** The distribution of V_run_ values. The peak at zero V_run_ here corresponds to the behavioural group A, whereas groups B and C form a single entity.

### Statistical analysis

The dataset was analyzed by ANOVA GLM. T_loc_, V_loc_, T_run_, V_run_, τ_ac_, and δ r were treated as dependent variables; group and sex as the categorical factors; body weights, litter index (taken as the inverse of the number of pups in the litter), and normalized number in order (taken as the ratio of the individual number in the order of testing to the number of pups in the litter) as the continuous predictors. Zero values for V_loc_ and V_run_ were excluded from the corresponding individual ANOVAs for V_loc_ and V_run_ so as to escape the zero-induced bias in mean values. Post-hoc Fisher LSD tests followed ANOVA if needed.

## RESULTS

The applied method of space potentials allowed a segmentation of the experimental trajectories into the localized walks and quasi-linear runs. The distinction was made based on a virtual space potential accumulated at each track point. The track fragments which were frequently revisited in their neighborhood became virtually “heated” and fell into the category of localized walks (see Appendix for the details). The smooth fragments left without re-visiting were considered as quasi-linear runs (Fig.2).

The blind open space trajectories consisted of localized walks, quasi-linear runs or their combination. The pups were assigned to one of the three groups (A, B, C) according to the individual values of T_loc_ as shown in Fig.3a (Table 1, see also the Methods above). The three groups corresponded to the visually observed locomotor strategies and trajectories’ types as follows: group A displayed preferentially localized walks (“drift”, the longest duration of T_loc_, 50< T_loc_ < 120 s); group B displayed a superposition of localized walks and quasi-linear runs (“drift and run”, 10s< T_loc_< 50s); the group C undertook almost immediate run trials (“run”, T_loc_< 10s). The pups of groups B and C reached the walls’ proximity and remained there till the end of their sessions. These ways of data clusterization produced the same groups of pups. The most objective way was to use the T_loc_ values, as shown in Fig.3a. The AC function calculated for the instant velocities of the localized walks showed a coherent decay during the first 2-3 AC times, paralleled by a very good match between the theoretical curve for Brownian motion and the two experimental ones (Fig.5a,b and Fig. 6a,b; red and blue markers for experimental data).

On a larger time scale the experimental mean squared displacement deviated significantly from the theoretically predicted linear trend (Fig.5a,b and Fig. 6a,b), being essentially sub-diffusive. It was paralleled by a prominent minimum in AC values, suggesting that frequent half-circle turns of the instant velocity vectors occured. Later, group B (blue markers) showed prominent oscillations of the AC function (Fig. 5), pointing to a looping pattern in the corresponding trajectories (Fig.2b). Group A behaved in a less regular way, with half-circles made but poorer completion of full circles (red lines closer to the abscissa than the blue ones, Fig.6a,b). For group B we compared the velocities of the localized walks and those of the quasi-linear run trials, because this group demonstrated both types of behaviour. Predictably, V_run_ was found to be greater than V_loc_ (p=0.002, Wilcoxson matched pairs test).

The groups A, B, and C differed significantly in quantitative locomotor parameters (Table 1). Briefly, group B shared the parameters of the localized walks with group A, and shared the parameters of quasi-linear runs with group C. V_loc_ did not differ between the groups A and B. Group C had a single pup which showed non-zero localized walk, so the group was excluded from the related ANOVA for parameters of localized walks (i.e. V_loc_, τ_ac_, δ r). The values of τ_ac_ and δ r did not differ between the groups B and C. Gender affected V_loc_ in a similar way in both behavioural groups where it was measured (B and C): V_loc_ was higher in females (1.3+0.1, N=23) than in males (0.9+0.1, N=19), F(_1,41_)6,7, p=0.014. T_loc_ differed between the groups, F (_2,50_)=192, p=0.000, thus validating its role as the discrimination criterion. T_run_ differed between the groups {F (_2,50_)=11, p=0.0001}, with group A showing the lowest values (p=0.00005 and p=0.0002 for the comparisons with groups B and C, according to Fisher LSD test). No difference in this parameter or a tendency to it was seen between groups B and C. V_run_ tended to differ between the groups {P=0.06{, due to this value for group A being lower than that for group B (p=0.03, Fisher LSD test). No difference or a tendency to it was seen for cohorts B and C. Body weight, litter index, or the normalized number in the order of testing were not significant continuous predictors for the locomotor parameters studied (with the exception of V_loc_, see above). Body weights of the pups didn’t differ significantly between the groups or sexes (24.3+1.4g and 22.1+1.1g for male and female pups, respectively; see Table 1 for the group difference). The same was seen for the litter index (7.6+0.5 and 8.0+0.4 for male and female pups, respectively; see Table 1 for the group differences) and the normalized number in the order of testing (6.6+0.1 for both sexes; see Table 1 for the group differences).

Body weights correlated negatively with the litter index (Rs=-0.68, p<0.05, N=51). Gender was not a predictor for the choice of the locomotor strategy. However, gender ratio differed between B and C groups (p=0.03, Fisher exact test). Namely, females prevailed in group B (10 out of 13 pups), whereas males prevailed in group C (7 out of 10 pups, Fisher exact test). Comparisons with A group (15 males, 13 females) did not reveal a significant difference in gender ratios.

## DISCUSSION AND CONCLUSIONS

The present study addresses the question of nonvisual navigation in immature rat pups. The rat, as crepuscular and nocturnal animals, must be able to navigate without significant visual flow [Hartman, 2011]. The development of locomotor abilities and appearance of the first nest-odour directed locomotor trials well before eye opening suggests that the basal levels of navigation in rats are nonvisual in their origins.

Indeed, we can see that the pups were able to follow at least two behavioural patterns (localized walks and quasi-linear runs), or combine them. Randomness was observed at the range of a few steps (see Fig.4 a,b and below) and perhaps might be expected further at the range of tens of minutes (see below).

**Fig.4.**
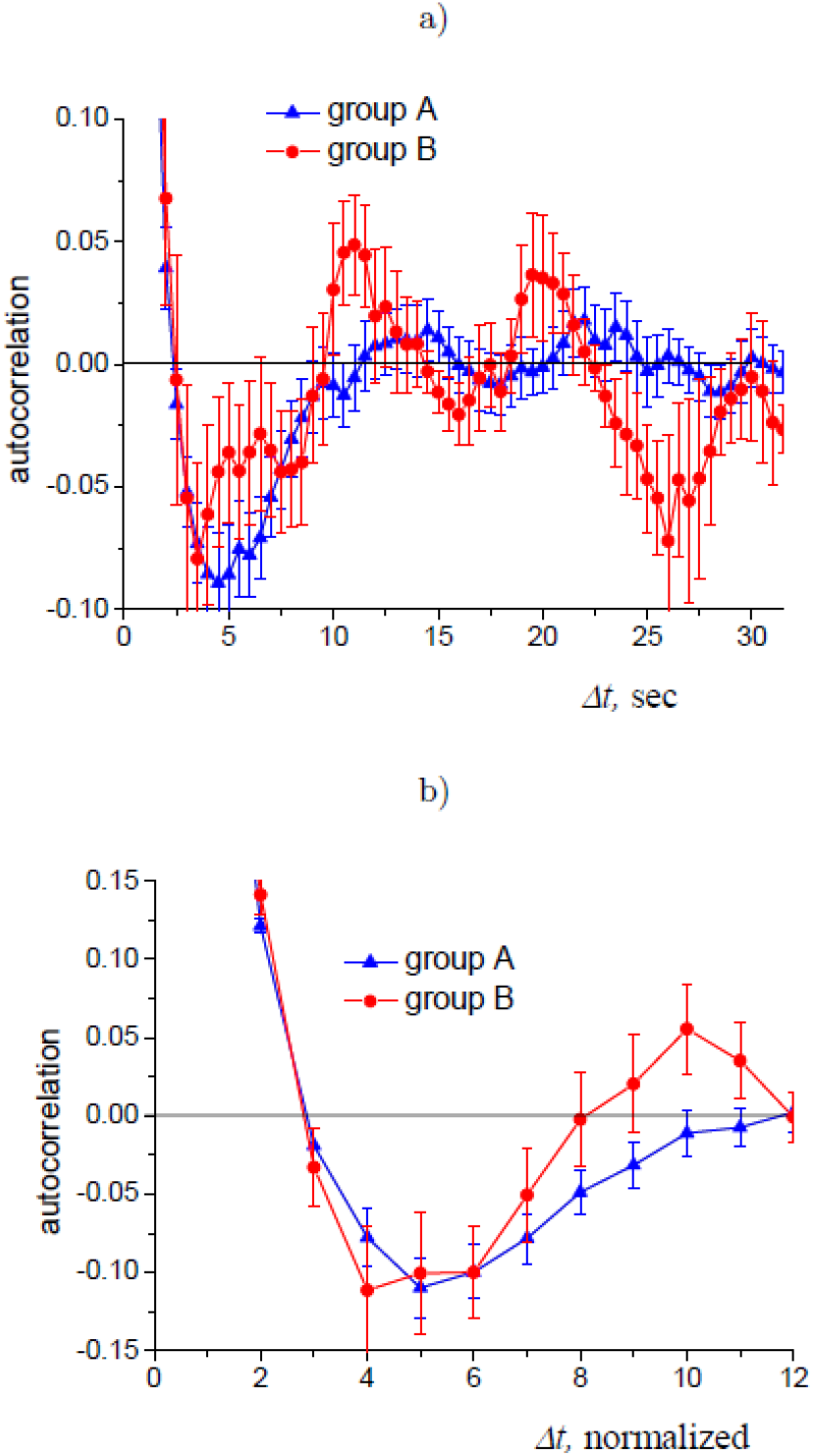
Velocity auto-correlation function averaged for groups A and B. **a)** averaging is done using auto-correlation as a function of time, **b)** averaging is done using auto-correlation as a function of time normalized via δ_ac_, see Appendix for more details. The error bars show the standard deviation of the means.

### Timescale of randomness in locomotion of previsual pups

According to the presented data, blind walks in normal previsual rat pups can be considered as an essentially random process at the scale of 2-3 seconds. It is deduced from a good match between the experimental data and the theoretical curves for Brownian walk (see Fig.4 a,b and Fig. 5 a,b).

**Fig. 5.**
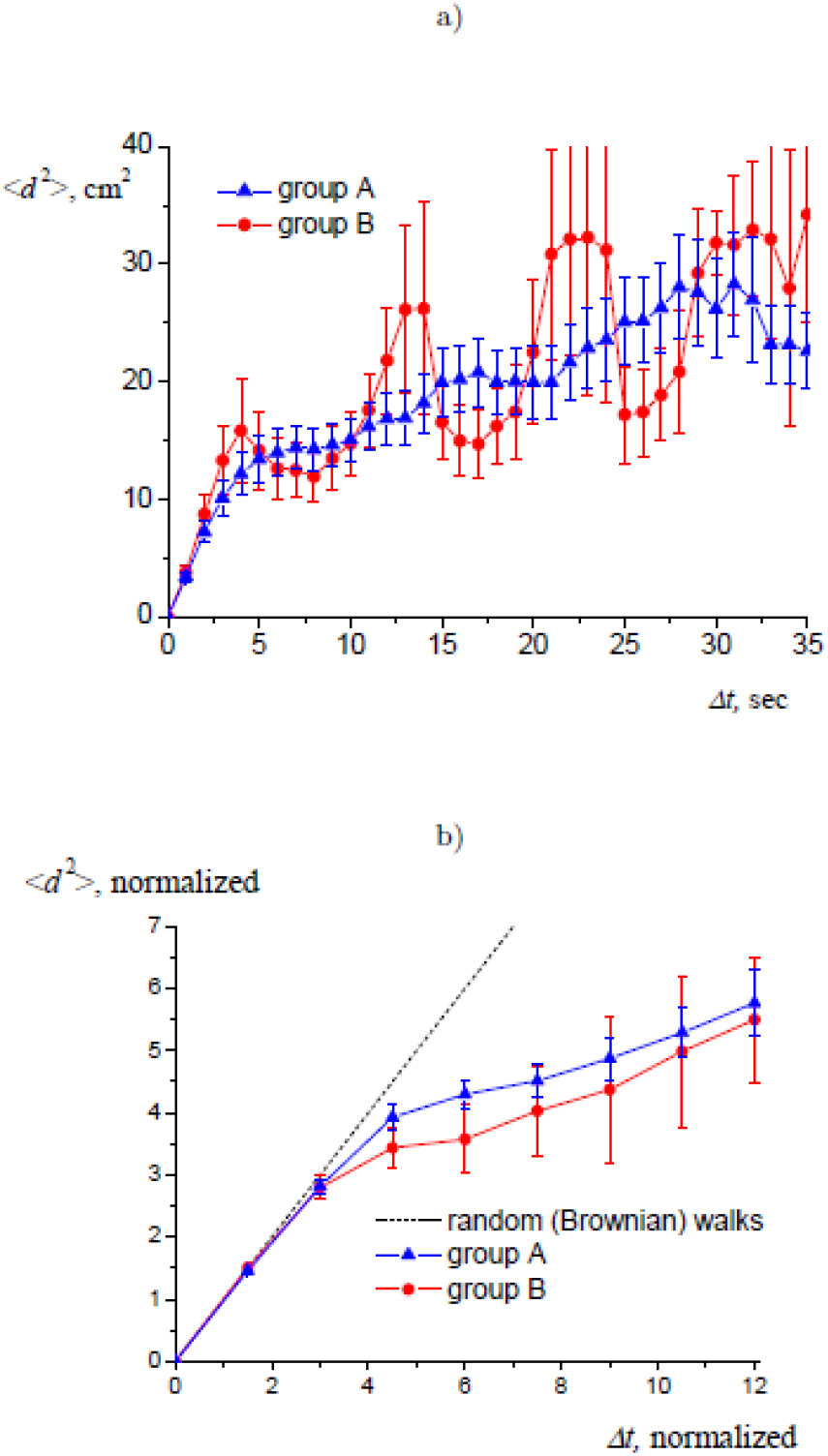
Mean squared displacement (MSD) for groups A and B. **a)** averaging is done using MSD as a function of time, **b)** averaging is done using MSD as a function of time normalized τ_ac_, see Appendix, Eq. (A6) and (A7) for more details. The error bars show the standard deviation of the mean. Dotted line in graph (b) shows the prediction of the Brownian motion model.

Namely, the mean squared displacement grows linearly and thus coincides with the theoretical curves for Brownian walks during the nearest 3-4 autocorrelation times (Fig.5 a,b; red markers for group A and blue markers for group B). It is further supported by the autocorrelation function calculated for the instant velocity vectors - it should decay exponentially for the Brownian processes, which can indeed be seen for the same range of 3-4 autocorrelation times (Fig.5 a,b; red markers for group A and blue markers for group B). The time scale implies that the locomotion is random at the range of a few steps.

### Localized walks are subdiffusive

On the time scale beyond the random domain the tracks can be viewed as a superposition of localized walks and quasi-linear runs. As mean body weights did not differ between the behavioural groups, we can conclude that the locomotor strategies did not depend directly on maturation. It means that “drifting” by localized walks is a behavioural strategy to choose, rather than a side-effect of reduced locomotor abilities.

Remarkably, the localized walks employ a mechanism of trajectories’ self-attraction. The mechanism works at the range of 5 seconds and more (see Fig.4 a,b). The experimental trajectories seen in the groups A and B displayed a revisiting effect, confirmed by a pronounced local minimum of the autocorrelation function (Fig.4 a,b) and experimental displacements lower than the Brownian one (blue and red experimental lines as compared to the black theoretical one (Fig.5 a,b). The first local minimum (which is 3-5 seconds, corresponding to 4-6 τ_ac_ on Fig.4 a,b) of the AC function strongly suggests that the pups turned back half-circle quite regularly. It is seems possible that the pups used their self-odour for navigation when lacking other sensory inputs. Olfactory sensing is the oldest navigation system, stemming from such evolutionarily old mechanisms as chemotaxis in invertebrates. Two odorants are sufficient for building a gradient space map [Jacobs, 2012]. In mammals navigating by olfaction, it is integrated and calibrated by sensory inflows from e.g. mechanosensory, auditory, and visual inputs [Jacobs and Schenk, 2003; Jacobs, 2012; Rossier and Schenk, 2003] to create parallel cognitive map [Jacobs, 2012].

Another possibility would be haptic space maps built via vibrissae sensing of the floor surface and/or instantly by tactile sensory inflow from the paws’ pads and tail. Although a smooth surface used in the present experiments (Fig.1) would not support the hypothesis of long-lived haptic space maps, additional experiments are needed to clarify the role of these possible references for blind locomotion.

Proprioceptive inputs from the paw pads and the tail would be able to support the trajectories re-attraction (Fig.2 a,b). Egocentric system of navigation would imply usage of such self-reference to minimize drift. On the other hand, minimizing the drift can suggest expectations of maternal retrieval. Rat mothers are known to quickly react on emission of isolation-induced 40kHz flat calls by retrieval of the separated infant [Hashimoto et al, 2001; Stern et al, 1984; Zeskind et al, 2011]. This would predict that the preference for localized walks coincides with a high rate of the isolation calls’ emission. This is to be clarified by the analysis of the USV calls emitted by the pups of groups A, B, and C (to be published).

To the best of our knowledge, the sub-diffusive (lower than Brownian) character of previsual locomotion has not been discussed in scientific literature yet. It would be interesting to investigate individual impacts of the factors (most probably, olfactory and proprioceptive) ensuring the tracks’ re-attraction.

If one considers the locomotor pattern as an intermittent process with trips and stops, then the localized walks are a prototype of dead reckoning during the extended stops. Dead reckoning (or path integration) is made by a navigator to deduce its location in space, with a reference to the starting point [Wallace et al, 2008]. The insufficiency of external stimuli in our experimental settings probably led to the prolongation and spatial extension of the dead reckoning phase (groups A and B).

### Quasi-linear raids as correlated random walks

The fast and smooth locomotion during the blind quasi-linear raids produced the impression that the locomotion was goal-directed. The raids ended in thigmotactic motion, which could be considered as an adaptive response. Thigmotaxis reduces the risk of potential predation, and thus increases security [Nemati et al, 2013; Whishaw et al, 2006].

However, the raids were undertaken in a random direction (see Supplement I) and missed features of an optimal escape run (were not necessarily directed to the nearest point of a wall), f.e see the track on Fig.2,b. Therefore, these fragments can be considered as a type of random walk. Since most animals, including the rat, tend to move “ahead”, i.e. mainly in the caudorostral direction, there is ‘persistence’ [Patlack, 1953; Codling et al, 2008] in their locomotor pattern. The random walks with increased persistence are correlated random walks [rev.in Codling et al, 2008]. Correlated blind walks should not have a directional bias, although they look more linear than a classical Brownian walk [Codling et al, 2008]. The presence of quasi-linear fragments and their random direction prompted us to hypothesize that the run trials are examples of correlated random walks. Another intriguing possibility is that the statistical law of run trials would be Levy flights, which are thought to be an energy-saving behavioural strategy in a search of rare goals [Benhamou, 2007; Edwards et al, 2007]. Levy flights are a realization of heavy-tailed distribution, i.e. very short and very long trips happen more frequently than it would be under Brownian law. Jump-like switch from localized walks to quasi-linear runs v_run_>v_loc_, see Results) would support the hypothesis. A larger statistics should bring clarity in the question, since correlated random walks involving a large number of steps produce Brownian displacement, whereas Levy flights result in super-diffusivity [Bartumeus et al, 2005].

In rat infants there is a mechanism to subserve directionality. The vital system of the directional sense is supported by a widespread network of functionally coupled head direction neurons [Dudchenko et al, 2019;Tan et al, 2017]. The system starts maturation well before eye opening in the rat [Tan et al, 2015; Tan et al, 2017] and will be finally tuned after the eyes’ opening [Basset et al, 2018; Tan et al, 2015]. Indeed, the ability to keep a direction of movement is already present by PND13 in previsual rat pups (groups B and C on Fig.2 b, c). Previously, pups of comparable age (PND14 and older), were reported to move along convoluted trajectories punctuated by stops while exploring an open field arena. The returns to a home-base were quasi-linear and faster than the outwards trips [Loewen et al, 2005]. In our experiment, the relation is reversed: outwards trips are faster and more linear. Punctuations by stops were seen here as a prolonged process (localized walks) with its internal dynamical features (see above). Quasi-linear runs appeared as outwards trips. It looked like an emerging quality of the system, due to a jump in velocity v_run_>v_loc_, see the Result section) and a loss of circuitry (Fig. 2 b,c). The reversed relation of inward/outward locomotion is obviously due to a novelty stress produced in pups and the absence of a home base. Quick and quasi-linear outward locomotion can be considered as a primary escape reaction, to optimize security[Nemati et al, 2013; Whishaw et al, 2006].

Animals tend to display saltatory patterns of movements [O’Brien et al, 1990], with pauses punctuating the tracks. The pauses are used to rest, forage or make a decision about the next move [Christensen et al 2015; Hunt et al, 2016;Christensen et al, 2021]. The locomotor strategies of previsual pups in the present experiments looked like a superposition of simplified trips and extended stops.

The present paper discusses locomotor strategies observed in previsual rat pups placed in a novel environment. Blind walks of previsual rat pups are considered as a superposition of two modes, the localized walks and the quasi-linear runs. Ontogenetic flow is directed from the first mode to the last one (as A-B-C transition). The localized walks employ an unknown mechanism of trajectories’ revisiting, which minimize the drift from the landing point and cause sub-diffusive nature of the locomotor mode. The track return mechanism can be based on olfactory and/or haptic sensory inputs. It is expectable that the emotional state of a pup would also affect the choice of the locomotor mode. Further experiments should clarify the issue.

Principles of blind autonomous locomotion of animals which have been evolved and optimized in various species can be used for building algorithms for various robotic devices, from underwater and underground ones to spaceships’ equipment. We propose to study ontogeny of laboratory rodents for this purpose, as new insights are available in quite feasible experimental set-ups. The developed algorithm of automated trajectories’ segmentation (see Appendix) is proposed for a wide usage by scientific communities.

## Supporting information

The space potentials' method for trajectories' analysis

## REFERENCES

1. Bartumeus, F., da Luz, M. G. E., Viswanathan, G. M., & Catalan, J. (2005). Animal search strategies: a quantitative random-walk analysis. Ecology, 86(11), 3078–3087. https://doi.org/10.1890/04-1806

2. Bassett, J. P., Wills, T. J., & Cacucci, F. (2018). Self-organized attractor dynamics in the developing head direction circuit. Current Biology, 28(4), 609–615. https://doi.org/10.1016/j.cub.2018.01.010

3. Benhamou, S. (2007). How many animals really do the Lévy walk?. Ecology, 88(8), 1962–1969. https://doi: 10.1890/06-1769.1

4. Bjerknes, T. L., Langston, R. F., Kruge, I. U., Moser, E. I., & Moser, M. B. (2015). Coherence among head direction cells before eye opening in rat pups. Current Biology, 25(1), 103–108. https://doi.org/10.1016/j.cub.2014.11.009.

5. Blekas, K., & Lagaris, I. E. (2007). Newtonian clustering: An approach based on molecular dynamics and global optimization. Pattern Recognition, 40(6), 1734–1744. https://doi: 10.1016/j.patcog.2006.07.012

6. Christensen, K., Cocconi, L., & Sendova-Franks, A. B. (2021). Animal intermittent locomotion: a null model for the probability of moving forward in bounded space. Journal of Theoretical Biology, 510, 110533. https://doi: 10.1016/j.jtbi.2020.110533

7. Christensen, K., Papavassiliou, D., De Figueiredo, A., Franks, N. R., & Sendova-Franks, A. B. (2015). Universality in ant behaviour. Journal of The Royal Society Interface, 12(102), 20140985. https://doi: 10.1098/rsif.2014.0985

8. Clarac, F., Brocard, F., & Vinay, L. (2004). The maturation of locomotor networks. Progress in brain research, 143, 57–66. https://doi.org/10.1016/S0079-6123(03)43006-9

9. Codling, E. A., Plank, M. J., & Benhamou, S. (2008). Random walk models in biology. Journal of the Royal society interface, 5(25), 813–834. https://doi: 10.1098/rsif.2008.0014

10. Dudchenko, P. A., Wood, E. R., & Smith, A. (2019). A new perspective on the head direction cell system and spatial behavior. Neuroscience & Biobehavioral Reviews, 105, 24–33. https://doi: 10.1016/j.neubiorev.2019.06.036

11. Edwards, A. M., Phillips, R. A., Watkins, N. W., Freeman, M. P., Murphy, E. J., Afanasyev, V.,… & Viswanathan, G. M. (2007). Revisiting Lévy flight search patterns of wandering albatrosses, bumblebees and deer. Nature, 449(7165), 1044–1048. https://doi.org/10.1038/nature09116

12. Jacobs, L. F., & Schenk, F. (2003). Unpacking the cognitive map: the parallel map theory of hippocampal function. Psychological review, 110(2), 285. https://doi: 10.1037/0033-295x.110.2.285.

13. Jacobs, L. F. (2012). From chemotaxis to the cognitive map: the function of olfaction. Proceedings of the National Academy of Sciences, 109(Supplement 1), 10693–10700. https://doi: 10.1073/pnas.1201880109

14. Jamon, M., & Clarac, F. (1998). Early walking in the neonatal rat: a kinematic study. Behavioral neuroscience, 112(5), 1218. https://psycnet.apa.org/doi/10.1037/0735-7044.112.5.1218

15. Hashimoto, H., Saito, T. R., Furudate, S. I., & Takahashi, K. W. (2001). Prolactin levels and maternal behavior induced by ultrasonic vocalizations of the rat pup. Experimental Animals, 50(4), 307–312. https://doi: 10.1538/expanim.50.307. PMID: 11515093.

16. Harshman, R. A. (1970). Foundations of the PARAFAC procedure: Models and conditions for an” explanatory” multimodal factor analysis. https://doi: 10.1016/j.patcog.2005.01.025.

17. Hartmann, M. J. (2011). A night in the life of a rat: vibrissal mechanics and tactile exploration. Annals of the New York Academy of Sciences, 1225(1), 110–118. https://doi: 10.1111/j.1749-6632.2011.06007

18. Hunt, E. R., Baddeley, R. J., Worley, A., Sendova-Franks, A. B., & Franks, N. R. (2016). Ants determine their next move at rest: motor planning and causality in complex systems. Royal Society open science, 3(1), 150534. https://doi: 10.1098/rsos.150534

19. Loewen, I., Wallace, D. G., & Whishaw, I. Q. (2005). The development of spatial capacity in piloting and dead reckoning by infant rats: use of the huddle as a home base for spatial navigation. Developmental Psychobiology: The Journal of the International Society for Developmental Psychobiology, 46(4), 350–361. https://doi: 10.1002/dev.20063

20. Nemati, F., Kolb, B., & Metz, G. A. (2013). Stress and risk avoidance by exploring rats: implications for stress management in fear-related behaviours. Behavioural processes, 94, 89–98. https://doi: 10.1016/j.beproc.2012.12.005.

21. O’Brien, W. J., Browman, H. I., & Evans, B. I. (1990). Search strategies of foraging animals. American Scientist, 78(2), 152–160. https://ui.adsabs.harvard.edu/abs/1990AmSci..78..152O

22. Patlak, C. S. (1953). Random walk with persistence and external bias. The bulletin of mathematical biophysics, 15(3), 311–338. https://doi: 10.1007/BF02476407.

23. Peyrache, A., Schieferstein, N., & Buzsaki, G. (2017). Transformation of the head-direction signal into a spatial code. Nature communications, 8(1), 1–9. https://doi: 10.1098/rstb.2013.0409.

24. Renner, M. J., & Pierre, P. J. (1998). Development of exploration and investigation in the Norway rat (Rattus norvegicus). The Journal of general psychology, 125(3), 270–291. https://DOI: 10.1080/00221309809595550.

25. Rossier J, Schenk F. Olfactory and/or visual cues for spatial navigation through ontogeny: olfactory cues enable the use of visual cues. Behav Neurosci. 2003 Jun;117(3):412–25. doi: 10.1037/0735-7044.117.3.412. PMID: 12802871. doi: 10.1037/0735-7044.117.3.412

26. Stern, J. M., Thomas, D. A., Rabii, J., & Barfield, R. J. (1984). Do pup ultrasonic cries provoke prolactin secretion in lactating rats?. Hormones and behavior, 18(1), 86–94. https://doi: 10.1016/0018-506x(84)90053-9. PMID: 6706322.

27. Tan, H. M., Wills, T. J., & Cacucci, F. (2017). The development of spatial and memory circuits in the rat. Wiley Interdisciplinary Reviews: Cognitive Science, 8(3), e1424.

28. Tan, H. M., Bassett, J. P., O’Keefe, J., Cacucci, F., & Wills, T. J. (2015). The development of the head direction system before eye opening in the rat. Current Biology, 25(4), 479–483. https://doi.org/10.1016/j.cub.2014.12.030.

29. Tocker, G., Borodach, E., Bjerknes, T. L., Moser, M. B., Moser, E. I., & Derdikman, D. (2018). Head-Direction drift in rat pups is consistent with an angular path-integration process. bioRxiv, 212852 https://doi.org/10.1101/212852.

30. Wallace, D. G., Martin, M. M., & Winter, S. S. (2008). Fractionating dead reckoning: role of the compass, odometer, logbook, and home base establishment in spatial orientation. Naturwissenschaften, 95(11), 1011–1026. https://doi: 10.1007/s00114-008-0410-z.

31. Whishaw, I. Q., Gharbawie, O. A., Clark, B. J., & Lehmann, H. (2006). The exploratory behavior of rats in an open environment optimizes security. Behavioural brain research, 171(2), 230–239. https://doi: 10.1016/j.bbr.2006.03.037.

32. Wills, T. J., Muessig, L., & Cacucci, F. (2014). The development of spatial behaviour and the hippocampal neural representation of space. Philosophical Transactions of the Royal Society B: Biological Sciences, 369(1635), 20130409. https://doi: 10.1098/rstb.2013.0409.

33. Zeskind, P. S., McMurray, M. S., Garber, K. A., Neuspiel, J. M., Cox, E. T., Grewen, K. M.,… & Johns, J. M. (2011). Development of translational methods in spectral analysis of human infant crying and rat pup ultrasonic vocalizations for early neurobehavioral assessment. Frontiers in psychiatry, 2, 56. https://doi: 10.3389/fpsyt.2011.00056

34. Zhao, Y., & Karypis, G. (2002, November). Evaluation of hierarchical clustering algorithms for document datasets. In Proceedings of the eleventh international conference on Information and knowledge management (pp. 515–524). https://doi: 10.1145/331499.331504

