## Supplementary material for "Locomotor strategies inside the blind walks: freely moving previsual rat pups in the open field test": The space potentials' method for trajectories' analysis

### APPENDIX: METHODS OF SPACE POTENTIALS APPLIED FOR TRAJECTORIES SEGMENTATIONS

#### A I. RAW TRACKS

Video of a pup's motion during 2 minutes is pre-processed by a special software which recognizes the pup's position. The output of the software is a set of $x$- and $y$-coordinates of the pup at the successive time instants. The temporal interval between them is $dt = 0.04$ second.

Fig. 2 in the main paper shows three examples of the pups' tracks. The track of the pup who has reached a wall without pronounced localized walks is shown in graph c); those shown in graphs a) and b) do contain such sections; however, the latter of these two tracks reaches the wall, whereas the former does not.

#### A II. REMOVING MINOR DISPLACEMENTS FROM THE TRACKS

There are fragments in the tracks where the displacements between the successive points are very small. Comparing the tracks and the video of the pup's motion we can see that these fragments correspond to time intervals when the pup actually does not move, but sits and (usually) washes itself. Small motions of the pup during washing lead to small displacements within these intervals. In our method we focus at the pup's walks, not at the above-mentioned minor motions. So we remove the points from the track if the displacement between the successive points is less than 0.3 cm (typical size of the pup's step).

#### A III. REMOVING TRACKS' FRAGMENTS AFTER REACHING THE VICINITY OF A WALL

A pup's motion usually changes after it reaches a wall: usually it runs along the wall and/or sits in the corner when it reaches it, see Fig2 b) and c) in the main paper. In our method we focus on the studies of the walks in the open field, so we remove the fragments of the tracks after the pup has reached the vicinity of the wall.

#### A IV. SELECTION OF THE LOCALIZED WALK FRAGMENTS OF THE TRACKS WITH A POTENTIAL-BASED METHOD

Points of the track after the above described pre-processing are denoted below as $x_k$, $y_k$, $t_k$; we also use the position of the k-th point $\mathbf{r}_k = \{x_k, y_k\}$ and the velocity at this point $\mathbf{v}_k = (\mathbf{r}_{k+1} - \mathbf{r}_{k-1})/(t_{k+1} - t_{k-1})$ . Note that the points are now *not* equidistant in time; this feature requires some peculiarities in defining the autocorrelation function which are discussed below.

The potential-based methods are routinely used for data clustering [1–3]. In our version of the method the $k$-th point of the track produces at a point $\mathbf{r}$ of the $(x, y)$ plane the potential:

$$p_k(\mathbf{r}) = \exp\left(-|\mathbf{r}_k - \mathbf{r}|^2/r_0^2\right), \tag{A1}$$

here $\mathbf{r}_k$ is the $k$-th point's position, $r_0$ is a constant. The parameter $r_0$ is chosen to be roughly equal to the pup's size, namely, $r_0 = 2.5$cm.

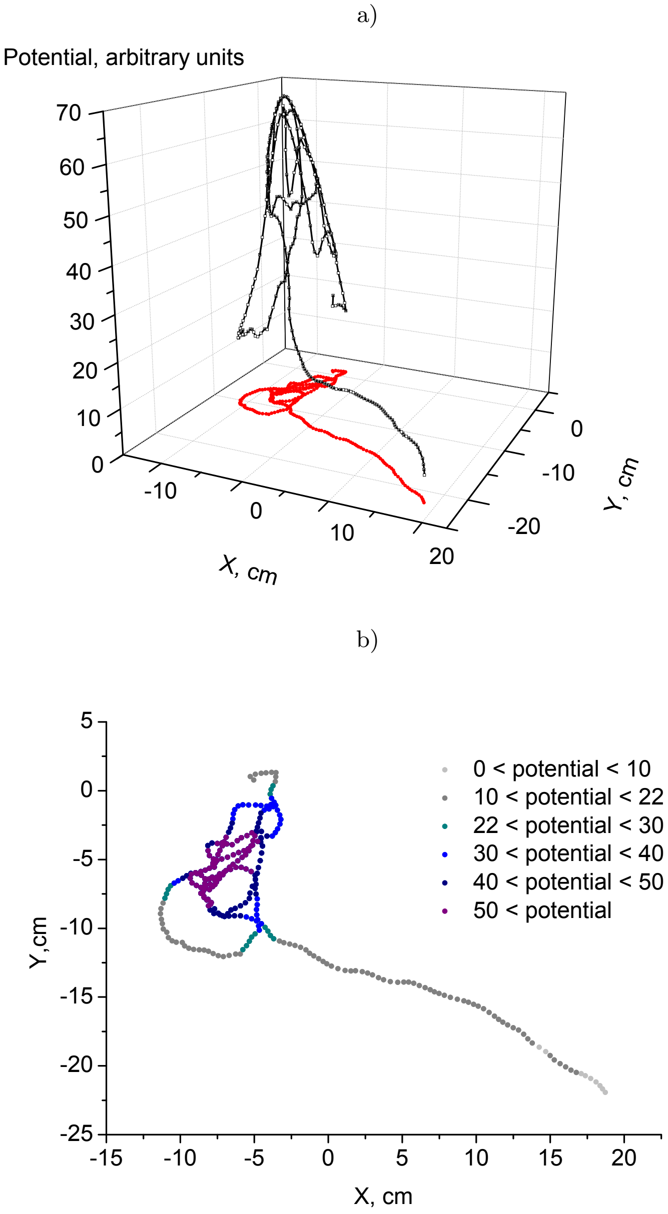


Figure A1. Illustration of the potential-based method used to separate the localized walk fragments of a track. Potential calculated via eq. (A2) is shown for each point of the track presented in Fig. 2b) of the main paper. Points where the potential is below threshold $P_{threshold} = 22$ are shown with gray circles in graph b), those with the potential above the threshold are shown with colored circles; these points correspond to the localized walk.

Total potential produced by all points $P(\mathbf{r}) = \sum_k p_k(\mathbf{r})$ is shown in Fig. 2 in the main paper.

The total potential at $j$-th point of a track is:

$$P_j = \sum_k p_k(\mathbf{r_j}) \tag{A2}$$

Note that we are using the short-range (namely, Gaussian) potential (A1) to be sure that the total potential of a cluster vanishes at a certain distance from the cluster even in the case of a cluster consisting of many points.

Figure A1 illustrates the potential at different points of the track shown in Fig. 2b) of the main paper (note that the part of the track near the wall has been already removed). One can see that the potential is higher at the 'entangled' fragments, corresponding to the localized walks. The points forming this fragment are those having the potential higher than a certain threshold. This threshold was chosen as $P_{threshold} = 22$.

Some tracks contain several entangled fragments. In these cases we select the longest one and consider it in the further analysis.

#### A V. VELOCITY AUTO-CORRELATION FUNCTION

The standard methods for calculating the autocorrelation function are derived for a time series with equal time intervals between the consecutive points. However, the series $t_k, \mathbf{v}_k$ is not equidistant in time, see section AII. Basing on this series we 'construct' a new equidistant time series $\tilde{t}_m, \tilde{\mathbf{v}}_m$ using a simple interpolation. Namely $\tilde{t}_m \equiv t_0 + m\,dt$ and $\tilde{\mathbf{v}}_m \equiv \mathbf{v}_k$, where $k$ is the number of the points closest in time to the initial series (i.e. $t_k$ is the instant most close to $\tilde{t}_m$).

Velocity autocorrelation function for each track is calculated as:

$$A(\Delta t) = \frac{\sum_m \tilde{\mathbf{v}}_m \tilde{\mathbf{v}}_{m+n}}{\sum_m |\tilde{\mathbf{v}}_m|^2} \tag{A3}$$

where $n = 0, 1, 2...$, $\Delta t = n\,dt$, $\tilde{\mathbf{v}}_m \tilde{\mathbf{v}}_{m+n}$ is a scalar product of the velocity vectors.

Fig. A2 shows the velocity autocorrelation as a function of the time delay $\Delta t$. Using an exponential fit of the autocorrelation function, we find the autocorrelation time $\tau_{ac}$ as a time where the fit is equal to $\exp(-1)$. Note that the found autocorrelation times are much longer than the initial temporal step of the tracking $dt$: $dt \ll \tau_{ac}$

We can see in Fig. A2 that successive points in the track are highly correlated, but then the correlation decreases. However, it does not vanish completely. To study this in more detail we calculate the mean displacement as a function of time as described in the next section.

#### A VI. MEAN SQUARED DISPLACEMENT (MSD) AS A FUNCTION OF TIME DELAY

To further study the randomness of the pups' motion we find the displacement $d_{j,k} = |\mathbf{r}_k - \mathbf{r}_j|$ between all pairs of points $j$ and $k$ and calculate the mean squared displacement (MSD) as a function of the time delay between the points:

$$< d^2 > (\Delta t) = \frac{\sum_{j,k:(n-0.5)\delta t < t_k - t_j < (n+0.5)\delta t} d^2_{j,k}}{\sum_{j,k:(n-0.5)\delta t < t_k - t_j < (n+0.5)\delta t} 1} \tag{A4}$$

where $\Delta t = n\,\delta t$, $n = 1, 2, 3...$, $\delta t$ is a time period which is much less than the total duration of the track $T$: $\delta t \ll T$.

Fig.A3 shows the displacement found via eq. (A4) using $\delta t = 1$ second for the two tracks shown in Fig. 2 of the main paper.

If the motion is completely random the squared displacement grows linearly with time. This model is denoted sometimes as a 'walk of a drunk man'; it describes Brownian motion and many other processes in physics, biology,

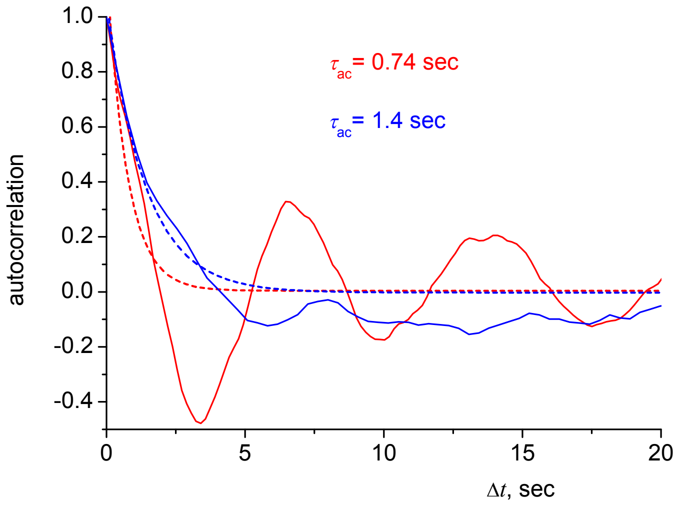


Figure A2. Velocity autocorrelation functions for localized walk fragments of the tracks shown in Fig.2 a) (blue) and b) (red) of the main paper. Dotted lines show exponential fits, the corresponding autocorrelation times are shown in the graph.

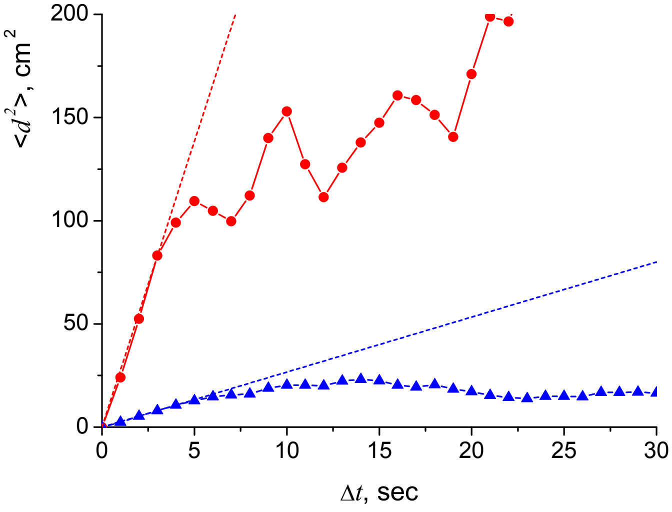


Figure A3. The mean square of the displacement between pairs of points delayed by time $\Delta t$ from each other calculated using eq. (A4) for the tracks presented in Fig. 2a) (blue) and b) (red) of the main paper. Dotted line shows the prediction of the Brownian motion model for each pup.

etc., see [4] and references therein. In Fig. A3 one can see that the squared displacement first grows linearly with time; in the graph we present the linear fits based on the first 4 seconds of the motion. However, for longer time intervals the displacement saturates. This means that displacements during short times are random, but the walk is localized if considered for a longer time period.

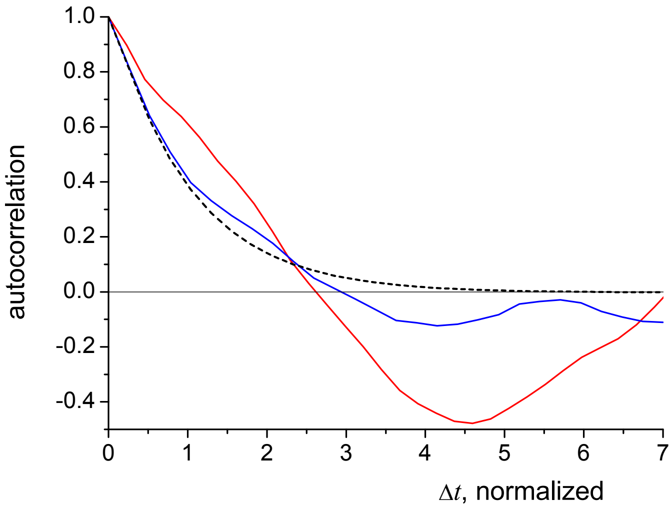


Figure A4. Velocity autocorrelation functions shown in Fig. A2 but time for each pup is normalized using its autocorrelation time $\tau_{ac}$. Dotted line shows the exponential fit which is common for both functions after the normalization of the time.

#### A VII. NORMALIZATION OF THE AUTOCORRELATION FUNCTION AND MEAN SQUARED DISPLACEMENT

Fig. A2 shows that the autocorrelation times are different for different animals, and in Fig A3 we can see that the rate of MSD increase at the stage of random motion is also different. Some aspects of averaging over different animals can be more conclusive if the averaging is done after the normalization of results described in this section.

Fig. A4 shows the autocorrelation functions as a function of normalized time delay

$$\Delta t_{norm} = \Delta t / \tau_{ac}.$$

Note that $\tau_{ac}$ is different for different pups. So the exponential fit is the same for both autocorrelation functions. One can see that this time normalization shows the common feature of the two autocorrelation functions: some anti-correlation within $3 \div 7\, \tau_{ac}$. More results are discussed in the main paper.

We normalize the mean squared displacement $< d^2(\Delta t) >$ using its value at the time equal to the autocorrelation time. The latter displacement is calculated via the following equation which is similar to (A4):

$$\delta r^2 = \frac{\sum_{j,k:0.8\tau_{ac}<t_k-t_j<1.2\tau_{ac}} d_{j,k}^2}{\sum_{j,k:0.8\tau_{ac}<t_k-t_j<1.2\tau_{ac}} 1} \tag{A5}$$

Note that within the Brownian motion model, $\delta r$ can be understood as the 'mean free path' of the pups' motion.

Calculating normalized mean squared displacement, the time delay $\Delta t$ is normalized using $\tau_{ac}$ for the pup in question, and the displacement is normalized using $\delta r$ for it. In more details, first, we calculate the autocorrelation time $\tau_{ac}$ for this pup. Then we calculate $\delta r^2$ for the pup using eq.(A5) and the mean squared displacement $< d^2 >$ using eq. (A4). The normalized mean squared displacement is:

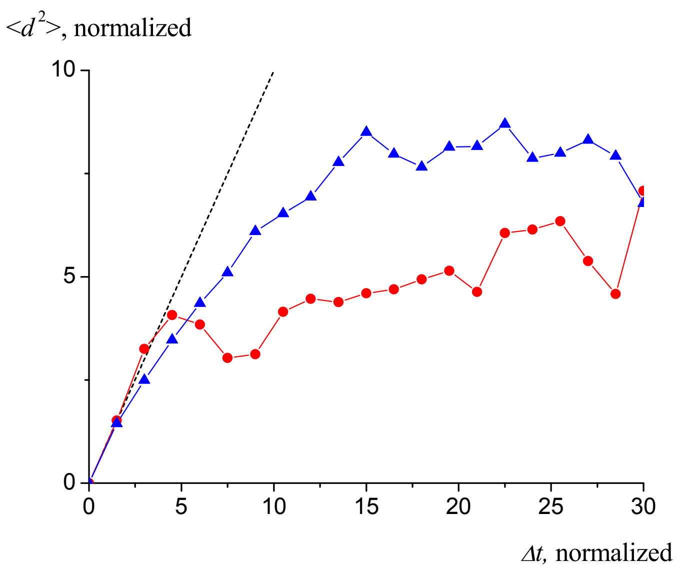


Figure A5. Same results as in Fig. A3 but time and displacement are normalized for each pup using its autocorrelation time $\tau_{ac}$ and the 'mean free path' $\delta r$. Dotted line shows the prediction of the Brownian motion model which is common for both functions after the normalization of time and displacement.

$$< d^2 >_{norm} (\Delta t_{norm}) = \frac{< d^2 > (\Delta t_{norm} \tau_{ac})}{\delta r^2} \tag{A6}$$

Fig. A5 shows the mean square of the displacement as a function of time delay. We chose $\delta t = 1.5\tau_{ac}$ for each pup so that the points $j$ and $k$ are delayed by time interval no less than ca. autocorrelation time even within the first bin (n=1 in eq. (A4)).

As we have already mentioned, the squared displacement grows linearly with time for a completely random motion. Moreover, for the normalization of time and displacement used the slope of the line is unity irrespective of $\tau_{ac}$ and $\delta r^2$. Thus, the displacement expected for a completely random walk is presented by a straight line shown as the dotted black line in Fig. A5. Thus, the used normalization allows comparing the results for different pups with the random walk model, as well as averaging the results for different groups of pups.

#### A VIII. AVERAGING OVER ANIMALS

Discussing the results for different pups we shall use a new index, $^{(p)}$ to denote the p-th pup.

The squared mean displacement averaged over a group of pups is:

$$\overline{< d^2 >_{norm}}(\Delta t_{norm}) = \sum_{p=1}^{p_{max}(\Delta t_{norm})} < d^2 >^{(p)}_{norm} / p_{max}(\Delta t_{norm}), \tag{A7}$$

where $p_{max}(\Delta t_{norm})$ is the number of pups in the group. The dependence of this number on $\Delta t_{norm}$ is due to the difference in the duration of the localized walks in the traces of different pups: for the shortest $\Delta t_{norm}$ the number

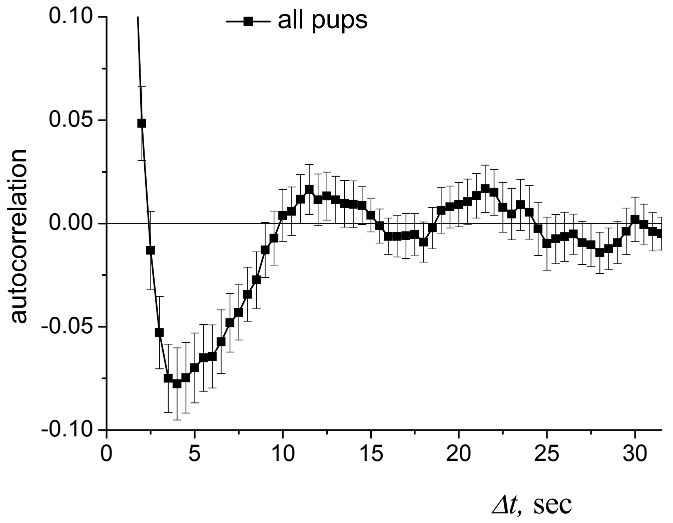


Figure A6. Auto-correlation function averaged for all pups. The error bars show the standard deviation of the mean.

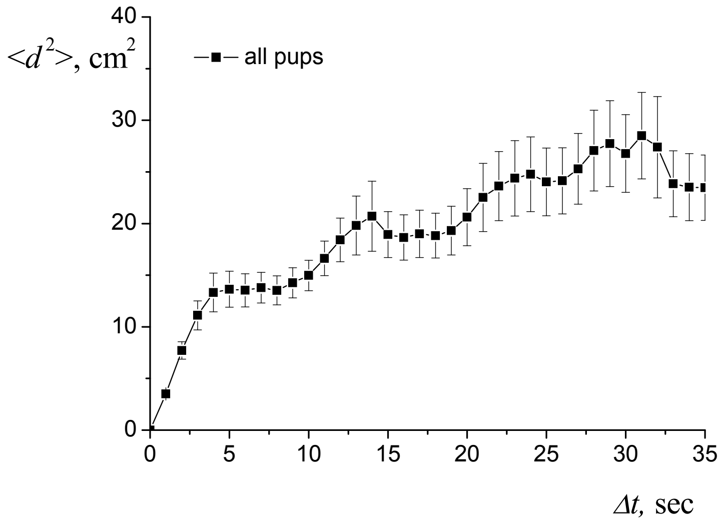


Figure A7. Mean squared displacement averaged for all pups. The error bars show the standard deviation of the mean.

$p_{max}$ is close to the total number of pups that demonstrated some localized walk fragments, whereas for longer $\Delta t_{norm}$ this number decreases.

The averaging of the autocorrelation functions are performed in a similar way.

The autocorrelation function and the mean squared displacement averaged over all pups are shown in Fig. A6 and Fig. A7, respectively.

[1] Warren Liao, T., Clustering of time series data - A survey, (2005) Pattern Recognition, 38 (11), pp. 1857-1874, www.elsevier.com/inca/publications/store/3/2/8/, doi: 10.1016/j.patcog.2005.01.025.

[2] Jain, A.K., Murty, M.N., Flynn, P.J., Data clustering: A review, (1999) ACM Computing Surveys, 31 (3), pp. 264-323, doi: 10.1145/331499.331504

[3] Blekas, K., Lagaris, I.E., Newtonian clustering: An approach based on molecular dynamics and global optimization, (2007) Pattern Recognition, 40 (6), pp. 1734-1744, doi: 10.1016/j.patcog.2006.07.012

[4] Codling EA, Plank MJ, Benhamou S. Random walk models in biology, (2008) J R Soc Interface, 5(25), pp. 813-834, doi: 10.1098/rsif.2008.0014. PMID: 18426776; PMCID: PMC2504494.
